# Morphological profiling of human T and NK lymphocytes identifies actin-mediated control of the immunological synapse

**DOI:** 10.1101/2020.01.17.910091

**Authors:** Yolla German, Loan Vulliard, Aude Rubio, Kaan Boztug, Audrey Ferrand, Jörg Menche, Loïc Dupré

## Abstract

The detection and neutralization of infected cells and tumors by cytotoxic lymphocytes is a vital immune defense mechanism. The immunological synapse orchestrates the target recognition process and the subsequent cytotoxic activity. Here, we present an integrated experimental and computational strategy to systematically characterize the morphological properties of the immunological synapse of human cytotoxic lymphocytes. Our approach combines high-content imaging with an unbiased, data-driven identification of high-resolution morphological profiles. Such profiling discriminates with high accuracy immunological synapse perturbations induced by an array of actin drugs in both model cell lines and primary lymphocytes. It reveals inter-individual heterogeneity in lymphocyte morphological traits. Furthermore, it uncovers immunological synapse alterations in functionally defective CD8^+^ T cells from immunodeficient patients carrying *ARPC1B* mutations. Our study thus provides a foundation for the application of morphological profiling as a powerful and scalable approach to monitor lymphocyte activation status in experimental and disease settings.

## Introduction

The immunological synapse (IS) is a complex cellular structure that sets lymphocyte activation and function during encounter with antigen-presenting cells and target cells. The canonical IS is characterized by a symmetrical architecture consisting of concentric rings of F-actin and integrins, while the antigen receptors occupy a central position^1, 2^. The lymphocyte spreading associated with IS assembly, as well as the molecular organization defining IS architecture, rely on actin cytoskeleton dynamics. In cytotoxic lymphocytes, including CD8+ T cells and NK cells, the IS is particularly important because it sustains the polarized delivery of cytolytic molecules such as perforin and granzymes towards target cells^3^. Indeed, activation of the integrin LFA-1 via an inside-out signaling from the T-cell receptor (TCR) in T cells, and several stimulatory receptors in NK cells^4–7^, leads to the formation of a tight adhesive ring allowing confinement of the degranulation process. Additional layers of control of lytic granule delivery at the IS are their polarization via the orientation of the microtubule organizing center ^8, 9^ and their restricted passage through pervasive actin cytoskeleton clearances^10^. Given the key events occurring at the IS, this structure is a window of choice to monitor lymphocyte activation and function. Indeed, the positioning and dynamic behavior of multiple receptors and signaling molecules have been characterized within the IS^11^, and alterations of the its architecture have been reported in multiple disease settings^12, 13^. However, the various microscopy approaches employed so far to characterize spatial organization of the IS have remained low throughput and have been restricted to the analysis of a limited number of morphological features. A more systematic in-depth assessment of the IS would better exploit this structure as a pivotal read-out for the characterization of lymphocyte activation and function.

Recent advances in high content imaging (HCI) now allow for the profiling of cells at a much richer level of detail and in an unbiased fashion. It has therefore been widely employed in cancer and toxicology research, in particular for screening drug effects on adherent cell lines and implementing genetic screens based on the siRNA, shRNA and CRISPR technologies^14–17^. However, HCI has not yet been applied to the study of leukocytes because of the difficulty to overcome the relatively poor adherence of these cells.

In this study, we report the implementation of an HCI approach that allows the high-resolution confocal imaging of T and NK cells stimulated over 2D surfaces functionalized with ICAM-1 and stimulatory antibodies, and the effect of pharmaceutical and genetic perturbations on the IS morphology. In addition to extracting a previously studied features related to staining of F-actin, LFA-1 and perforin, we develop an unbiased analytical approach allowing high-dimensional profiling and clustering of IS morphologies. Our data shows non-identical perturbations caused by drugs affecting different facets of actin cytoskeleton remodeling and highlights that actin cytoskeleton integrity is required not only for lymphocyte spreading but also for lytic granule polarization and LFA-1 distribution. Application of our HCI pipeline to lymphocytes isolated from human blood reveals distinct morphological profiles in individual healthy donors. Furthermore, our method allows characterizing synapse defects in untransformed CD8^+^ T cells from ARPC1B-deficient patients, illustrating its potential to identify disease-related synapse alterations and to predict functional defects, such as cytotoxicity.

## Results

### Morphological profiles of T cell and NK cell immunological synapses

In order to systematically analyze the morphological profile of lymphocyte populations, we here sought to develop an adapted HCI workflow. It consisted in seeding cells of interest on stimulatory surfaces in microwells of 96- or 384-well plates, fixation and staining with combinations of fluorescent dyes and antibodies. Confocal images were acquired on an Opera Phenix high-content screening system and analyzed with CellProfiler^18^ to automatically segment individual cells and extract features pertaining to cell morphology and each of the fluorescent markers (Fig. 1a). As proof of concept, we first applied our approach to NK-92 and Jurkat cells, two human cell lines commonly used as models for NK cells and T cells, respectively. Cell morphologies were compared upon interaction with a neutral poly-L-lysine (PLL) surface or co-stimulation with the LFA-1 ligand ICAM-1 and stimulatory antibodies (Ab) in order to evoke IS assembly. Upon co-stimulation with ICAM-1 and anti-NKp30 / NKp46 Ab, NK-92 cells spread, emitted F-actin-rich peripheral pseudopodia and polarized perforin-containing granules towards the center of the cell to substrate interface (Fig. 1b and **Fig. S1a**). These observations are in line with the characteristics of the IS from cytotoxic lymphocytes^11, 19^, therefore validating our high-throughput stimulation and staining procedure. Based on literature describing the IS and reporting a polarization of F-actin and lytic granules in NK cells^20–22^, we first selected quantitative features pertaining to the F-actin and perforin stainings and extracted them as mean values per field of view averaged across 3 experiments. We also included features related to the nucleus, available since the DAPI staining was used in a primary nucleus segmentation step before the identification of the cytoplasms around the nuclei (Fig. 1c and **Table S1**). Increase in F-actin intensity and cell area were prominent features of the stimulation, as compared to the PLL condition. Furthermore, the number of perforin-containing granules detected at the cell to substrate interface increased upon stimulation, which is indicative of their polarization towards the IS. Interestingly, this polarization process was associated with a relative spreading of the area covered by lytic granules, supporting the notion of multiple docking domains at the synapse^23^. Our analysis also highlights that increase of nucleus area is a typical feature of the IS in the NK-92 cells. Interestingly, nucleus area appears to increase along with F-actin intensity when assessed across 3 experiments (Fig. 1d), suggesting that nucleus flattening and F-actin polymerization are related events, probably as components of the cell spreading mechanism. Of note, the absolute values for F-actin intensity were higher in one of the 3 experiments, possibly resulting from differences in staining quality. This indicates that absolute value for staining intensities across experiments should be considered with caution. To further estimate morphological heterogeneity in individual cells, F-actin intensity was assessed on a per-cell basis (Fig. 1e), rather than on a per field of view basis. The unimodal increase of F-actin intensity driven by the stimulation of NK-92 cells indicates a relatively homogenous activation and IS assembly in these cells. It also validates our approach to consider mean cell measurements on a per field of view basis.

**Fig. 1.**
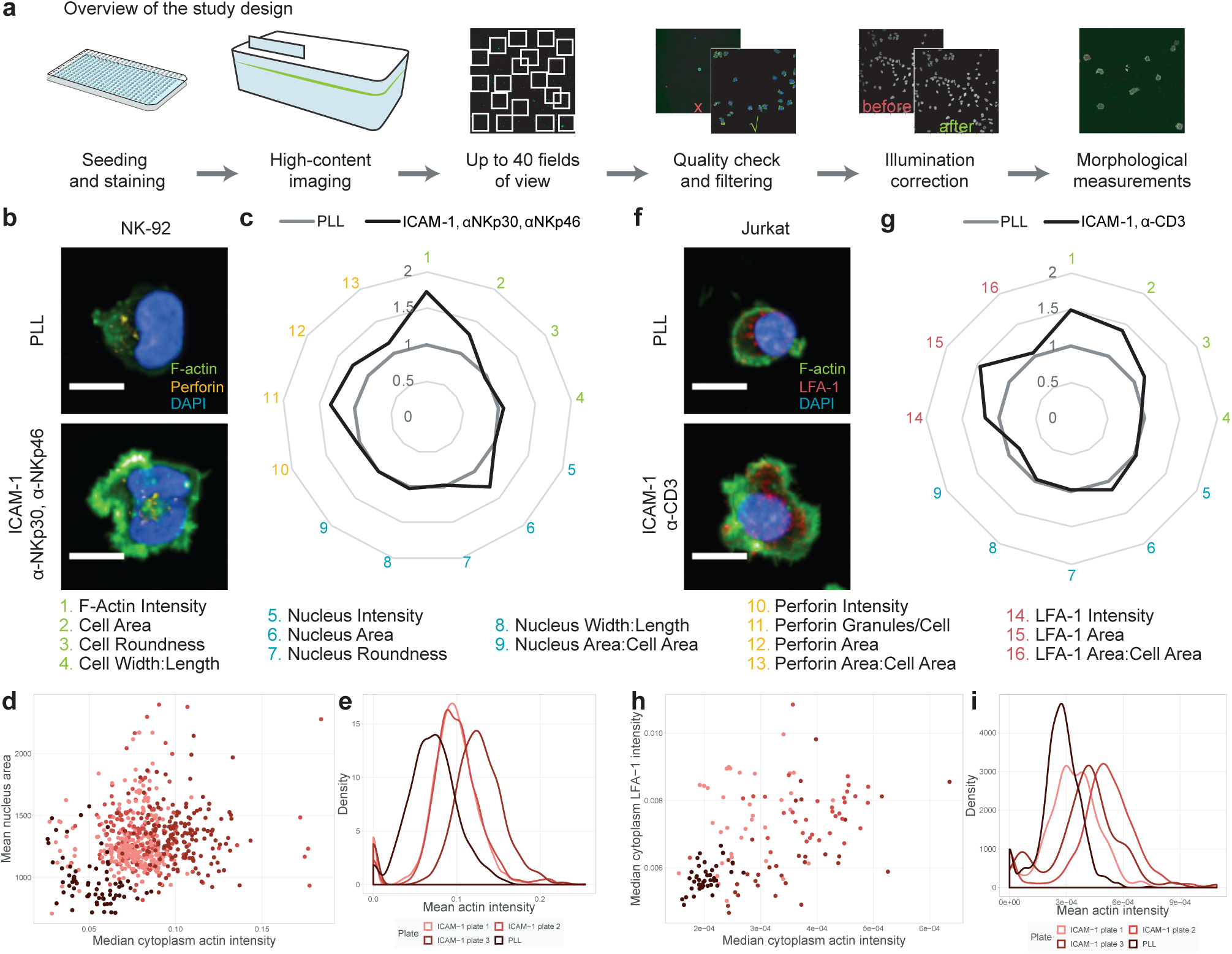
High content imaging of the immunological synapse in lymphocytic cell lines. **a.** Schematic representation of the High content imaging pipeline. **b.** Representative images of NK-92 cells seeded on Poly-L-Lysine (PLL) or ICAM-1 plus anti-NKp30 / NKp46 Ab, stained for F-actin (green), perforin (yellow) and the nucleus (DAPI). Scale bars: 10 µm. **c.** Selected IS features analysed as fold change of ICAM-1 plus anti-NKp30 / NKp46 Ab over PLL. The data represent the mean of three separate experiments (n= 933-5860 cells). **d.** Mean nucleus area in pixels and median F-actin intensity per image across PLL and ICAM-1 plus anti-NKp30 / NKp46 Ab conditions. **e.** F-actin intensity distribution per cell across PLL and ICAM-1 plus anti-NKp30 / NKp46 Ab conditions. **f.** Representative images of Jurkat cells on PLL or ICAM-1 plus anti-CD3 Ab, stained for F-actin (green), LFA-1 (red) and the nucleus (DAPI). Scale bars: 10 µm. **g.** Selected IS features analysed as fold change of ICAM-1 plus anti-CD3 Ab over PLL. The data represent the mean of triplicates (n= 125-940 cells). **h.** Median F-actin and LFA-1 intensity per image across PLL and ICAM-1 plus anti-CD3 Ab conditions. **i.** F-actin intensity distribution per cell across PLL and ICAM-1 plus anti-CD3 Ab conditions.

We then applied our HCI workflow to Jurkat cells, which were co-stimulated with ICAM-1 and anti-CD3 Ab. We selected 12 features pertaining to the F-actin, LFA-1 and DAPI stainings to monitor hallmarks of the T cell IS^11^. As compared to the neutral PLL stimulation, LFA-1/CD3 co-stimulation led to cell spreading, assembly of a peripheral F-actin ring-like structure and redistribution of the integrin LFA-1 as an inner belt at the cell to substrate interface (Fig. 1f and **Fig. S1b**), which are characteristic for the IS^11, 24^. Our quantification over multiple fields showed that similarly to NK-92 cells, F-actin intensity, cell area, LFA-1 intensity and LFA-1 area are prominent features of the Jurkat cell IS (Fig. 1g and **Table S2**). Likewise, F-actin and LFA-1 intensities correlated in individual fields of view with a Pearson correlation coefficient of 0.50 (Fig. 1h). At the single cell level, F-actin clearly increased in response to the ICAM-1 and anti-CD3 Ab stimulation, despite noticeable heterogeneity in both stimulated and unstimulated cells (Fig. 1i). When taken together, the data collected on the assembly of the IS in NK-92 and Jurkat cells highlight F-actin intensity rise and cell spreading as common traits. However, in line with the distinct appearance of their actin rich peripheral protrusions, Jurkat cells, but not NK-92 cells, became rounder upon activation. Furthermore, while NK-92 flattened their nucleus, the effect was not marked in Jurkat cells, indicating a distinct cell spreading behavior. Overall, our data validate the reliability and power of HCI with high spatial resolution to unbiasedly define the morphological profiles of lymphocytes in response to stimulatory regimens.

### Cytoskeleton drugs induce non-identical alterations of the NK cell IS

Given the prominent actin remodeling activity sustaining IS assembly, we next exploited our HCI approach to monitor how pharmacological modulation of cytoskeletal dynamics would affect IS architecture. NK-92 cells were treated with seven drugs known to target distinct aspects of actin and acto-myosin dynamics, which were used at 3 concentrations in order to detect possible dose-dependent effects. NK-92 cells were pre-treated with drugs for 30 minutes before seeding on ICAM-1 and anti-NKp30 / NKp46 Ab. In a first step, the effects of these drugs on the previously selected quantitative morphological features of the IS were assessed.

Upon treatment with Latrunculin B, which binds actin monomers and inhibits actin polymerization, a concentration-dependent decrease in F-actin intensity was detected as compared to the untreated control (Fig. 2a), in concordance with an expected reduction in actin polymerization. However, IS assembly was not fully impeded as revealed by a minor reduction of cell area. Remarkably, Latrunculin B treatment elicited an increase of the number of perforin granules and the area they occupied at the cell to substrate interface. This might reflect impaired exocytosis and accumulation of aberrantly spread lytic granules at the IS. Such explanation is in agreement with the role of actin dynamics in facilitating the docking and exocytosis of lytic granules^21, 25^. Upon treatment with Jasplakinolide, which stabilizes actin, a mild decrease in F-actin intensity was detected at 1 µM, supporting the notion that actin turnover is required to fuel polymerization^26, 27^(Fig. 2b). A higher concentration of 2.5 µM was tested but could not be exploited because of its apparent detrimental effect on cell viability. In comparison to Latrunculin B, Jasplakinolide treatment elicited an increase in the perforin-related features, confirming that actin turnover is required for lytic granule exocytosis^25, 27^. Treatment with the myosin inhibitor Blebbistatin induced a slight increase in F-actin intensity at 5 and 10 µM (Fig. 2c). More strikingly it increased the number of granules detected at the synapse and the area they occupied, consistent with previous findings that Blebbistatin hinders granule exocytosis without affecting their polarization^28, 29^. Treatment with the ROCK inhibitor Y-27632 affected F-actin intensity, cell area and lytic granule features similar to those elicited by Blebbistatin treatment (Fig. 2d), in agreement with the activity of ROCK as an upstream regulator of acto-myosin contractility. Upon CK-869 treatment, a concentration-dependent decrease in F-actin intensity was detected (Fig. 2e), showing, as expected, that ARP2/3 complex inhibition reduced actin polymerization^30^. Moreover, CK-869 treated cells displayed reduced radial spreading, as shown by decreased area and increased cell width to length ratio, indicative of a severe impairment of IS assembly. A distinct property of CK-869 treatment was a reduction in the number of and area covered by perforin granules, possibly reflecting the inability of CK-869 treated cells to polarize lytic granules towards the stimulatory surface because of defective IS assembly. Upon treatment with Wiskostatin, an inhibitor of WASP, which drives ARP2/3-dependent actin branching, a slight increase in F-actin intensity was measured for the two lowest concentrations (Fig. 2f). In comparison with CK-869, Wiskostatin displayed minor effects on perforin features. This suggests that the ARP2/3 activator WASP plays a limited role in the overall actin polymerization rate at the IS and in lytic granule polarization and secretion^24, 31^. Treatment with the pan-formin blocker SMIFH2 led to a concentration-dependent increase in F-actin intensity (Fig. 2g). Low-concentration SMIFH2 treatment resulted in an increase in perforin intensity and area. Collectively, these data indicate that drugs affecting different facets of actin remodeling differentially altered the assembly of the NK cell IS.

**Fig. 2.**
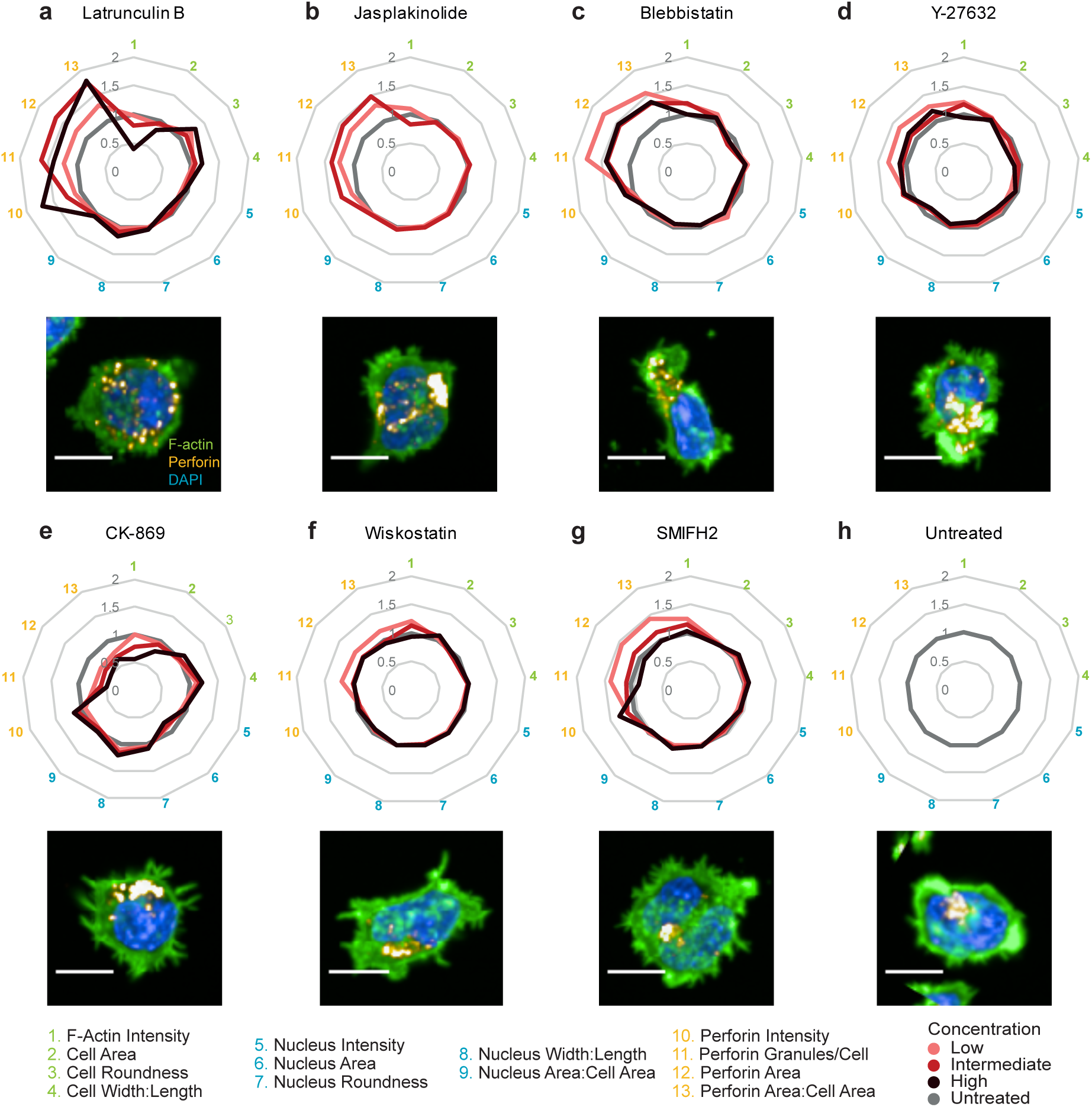
Drug treatments yield changes to immunological synapse morphology. Graphs representing the fold change of IS parameters and representative images of NK-92 cells seeded on ICAM-1, anti-NKp30 and anti-NKp46, stained for F-actin (green), perforin granules (yellow) and nuclei (DAPI) and treated with three concentrations of **a**. Latrunculin **b.** Jasplakinolide, **c.** Blebbistatin, **d.** Y-27632, **e.** CK-968, **f.** Wiskostatin, and **g.** SMIFH2 with respect to the untreated control. **h.** Untreated control. The data represent the mean of triplicates for each concentration (1425-5541 cells). Scale bars: 10 µm.

To further assess whether the morphological alterations inflicted by the drugs could be distinguished one from another, we trained a random forest classifier based on the 13 selected features. The image set was split to carry out a parameter optimization and to validate the performance of the model. The obtained overall accuracy of 69% and *F*_1_ score of 0.7 confirmed that our method performed relatively well at distinguishing the morphological effects of the actin drugs. The confusion matrix shows that most drugs were predicted with high accuracy based on the corresponding image features, while the morphological effects of Blebbistatin and Y-27632 could not be easily distinguished, in line with their highly related mechanism of action (**Fig. S2a**). It also confirmed F-actin intensity as a major discriminating feature and identified cell eccentricity and roundness as key features to account for the morphological alterations induced by the drugs (**Fig. S2b**).

Overall, our comparative image-based analysis of the effects of different drugs affecting actin and acto-myosin dynamics reveals that distinct effects on actin turnover and polymerization yield distinguishable IS morphologies but converge in affecting lytic granule positioning. This supports the notion that multiple actin-dependent steps control lytic granule docking and exocytosis^32^.

### High-resolution morphology profiling of NK cells upon drug treatment

To further enrich our morphological analysis of the IS in the context of drug treatment, we considered additional morphological features beyond the previously analyzed quantitative features. From 1898 measured features, a set of 383 features was retained following filtering of non-informative and redundant features. In order to visualize the information contained in this large feature set, as well as to quantify the significance of morphological changes upon drug treatment as compared to untreated cells, we applied a UMAP dimensionality reduction. This allows the visualization of all cell images, recapitulating in a ‘morphological space’ the relation between the morphology they display, by summarizing the variation of the 383 features into two dimensions (**Fig. S2c**). By examining the relation between these features, we saw that both the different types of measurements acquired and the different biological objects studied provided complementary and non-redundant information about the global changes occurring between images and between treatments (**Fig. S2d**). This also showed that none of these morphological features were repeating technical confounders, such as experimental plate position effect or cell count. As clearly visible in the morphological space, images of cells treated with Latrunculin B, Jasplakinolide and CK-869 clustered away from the untreated cells and from one another, most likely owing to these drugs having prominent and distinct effects on the ability of NK-92 cells to assemble the IS (Fig. 3a, b and e). In comparison, morphologies of cells treated with Blebbistatin, Y-27632, Wiskostatin and SMIFH2 appeared to be less distinct from the untreated condition and to cluster at close vicinity to one another (Fig. 3c, d, f and g). The three concentrations assayed per treatment fell into distinct sub-clusters, clearly indicating dose-dependent effects, as detailed for CK-869 and SMIFH2 (**Fig. S2e** and **f**). All drug-evoked morphological profiles were found to be significantly distant from the untreated state. Indeed, the median robust Mahalanobis distances between the fields of view per treatment and their matching negative controls are larger than expected at random (**Fig. S2g**)^33, 34^. To get insight into the nature of the changes that are causative of the observed clusters on the UMAP representation, we trained a random forest classifier on the set of 383 features. This achieved a satisfactory performance, as shown on the confusion matrix (Fig. 3h) with an *F*_1_ score and an accuracy of 0.89 and 89%, respectively. The importance of each feature for the classification was proxied by the average increase in accuracy obtained by including the given feature in a decision tree. In particular, our analysis shows that CK-869 treatment mostly affected nucleus and cytoplasm shape descriptors (Fig. 3i), while SMIFH2 treatment altered radial intensity distributions in the cytoplasm (Fig. 3j). Only four features described intensities in the cytoplasm within our feature set. Interestingly, those few features were in average increasing the model accuracy the most, strengthening the necessity, but not sufficiency, of actin intensity measurements to profile the IS. Features pertaining to lytic granules also played a determinant role in reinforcing model accuracy, providing further evidence of a tight regulation of lytic granule distribution at the IS. Our data therefore demonstrates the ability of the unbiased profiling to identify relevant spatially localized events and characterize perturbed cell states with high-resolution power.

**Fig. 3.**
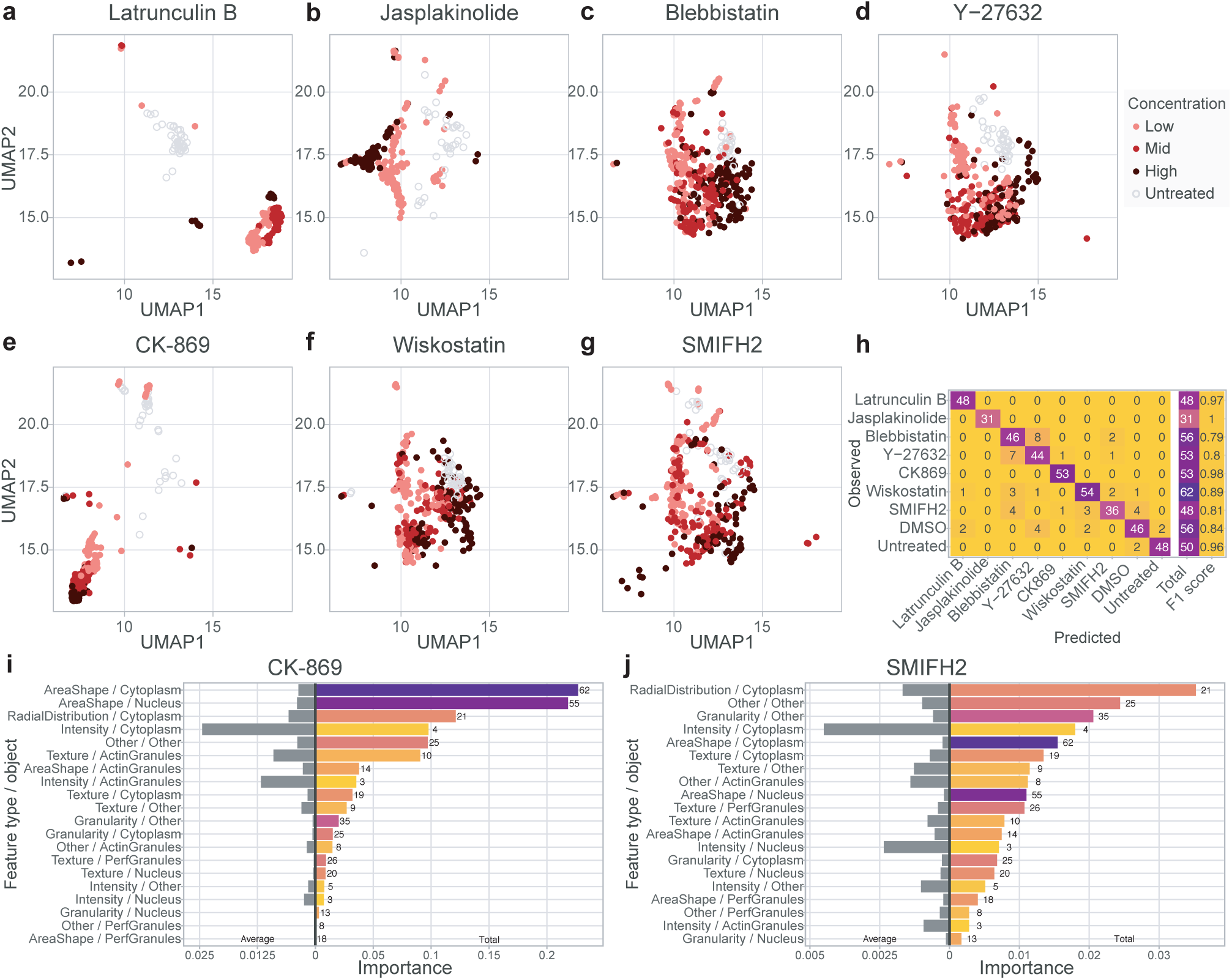
Morphological profiling of the NK cell immunological synapse upon drug treatment. Drug-treated NK-92 cell images were analysed with CellProfiler for an array of measurements and visualized using UMAP to position drug treated cells with respect to untreated cells from the same row. **a**. Latrunculin **b.** Jasplakinolide, **c.** Blebbistatin, **d.** Y-27632, **e.** CK-968, **f.** Wiskostatin, and **g.** SMIFH2. **h.** Confusion matrix and class-wise performance on held-out data of a random forest model trained to predict drug treatment based on the morphology of NK-92 cells seeded on ICAM-1, anti-NKp30 and anti-NKp46. **i-j** Total and average importance for the prediction of morphological features per measurement type and biological object described of NK-92 cells seeded on ICAM-1, anti-NKp30 and anti-NKp46 and treated with **i.** CK-869 or **j.** SMIFH2.

### Morphological profiling of primary human NK cells upon drug treatment

We next applied our HCI approach to assess the susceptibility of primary human lymphocytes from different donors to cytoskeletal drugs. For that purpose, NK cells were purified from the peripheral blood of three normal donors, treated with four concentrations of either CK-869 or SMIFH2, and stimulated with ICAM-1 and anti-NKp30 / NKp46 Ab (Fig. 4a). Although the untreated cells from the three different healthy donors displayed variation in morphology, an actin-rich IS with the lytic granules concentrated in one area towards the cell periphery was observed upon stimulation. The four tested concentrations of CK-869 caused a marked decrease in F-actin intensity in the NK cells from the three donors, demonstrating the capacity of our approach to detect actin cytoskeleton alterations in primary lymphocytes. Notably, the area covered by the perforin granules, taken as an absolute value or divided by the cell area, was increased in the CK-869 treated NK cells from the three donors, showing a clear dose-dependent response (Fig. 4b). This effect is opposite to what was measured in the NK-92 cells, highlighting contrasting responses of model cell lines and primary cells. Moreover, the four tested concentrations of SMIFH2 also caused a decrease in F-actin intensity in the NK cells from the three donors, thereby highlighting the importance of formins for actin remodeling at the IS (Fig. 4c). SMIFH2 treatment also strongly affected the distribution of perforin granules. Interestingly, in NK cells from donors 1 and 2, a dose dependent reduction of both perforin granule number and covered area was observed, a response opposite to that to CK-869. In contrast with donors 1 and 2, low concentrations of SMIFH2 resulted in an increase of perforin granule number and covered area in NK cells from donor 3. It should be noted that lower number of perforin granules were detected in the untreated cells from this donor, possibly influencing the response to the tested drugs. Those observations highlight the potential of HCI to identify features underlying inter-donor variability upon stimulation and treatment of lymphocytes populations, which may be explained by the phenotypic variation of each donor’s NK cells^35^. Together, the dataset collected on primary NK cells demonstrates that HCI is amenable to the morphological profiling of primary human lymphocytes in the context of drug treatments and that it can discriminate specific responses from individual donors.

**Fig. 4.**
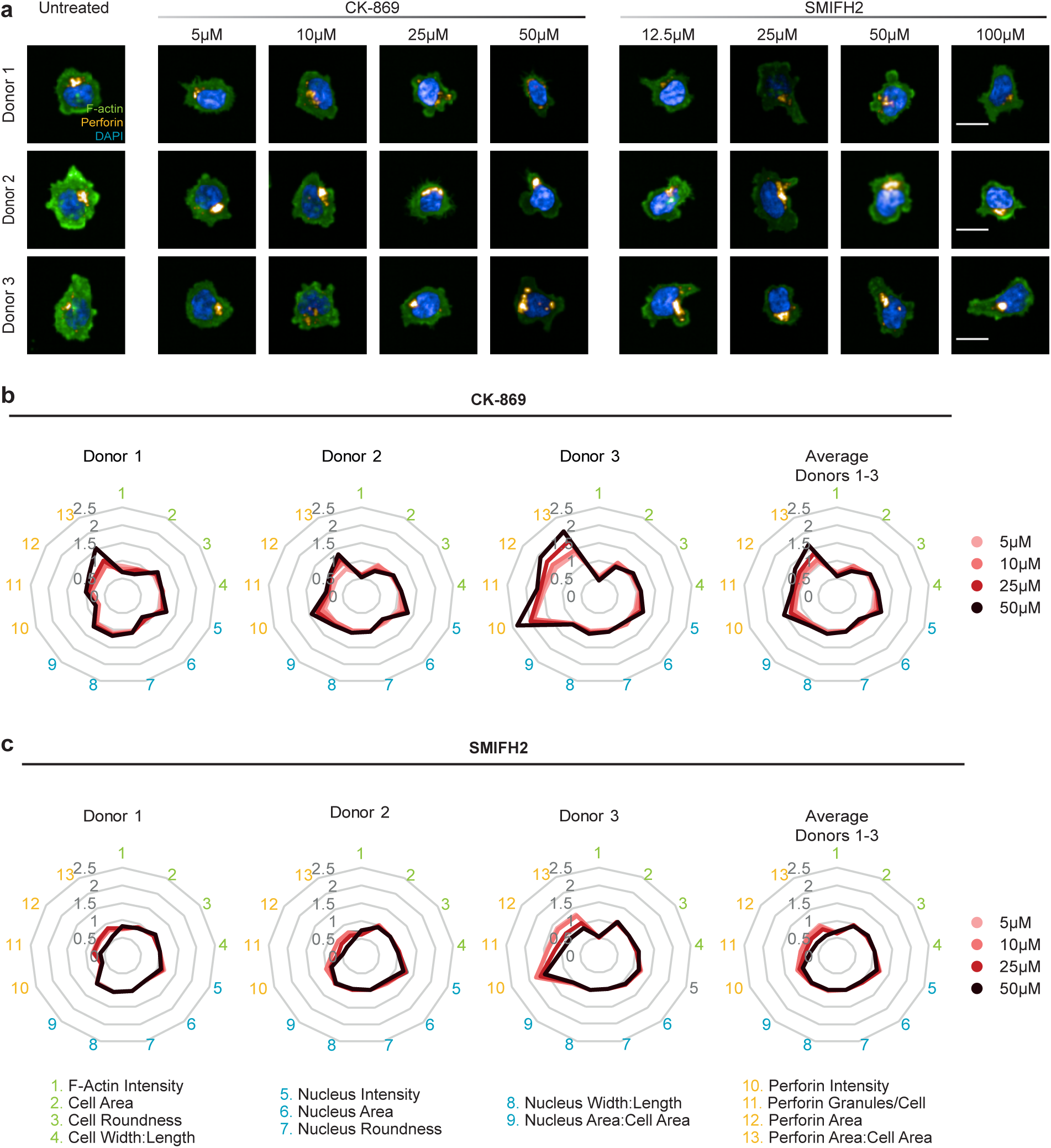
CK-869 and SMIFH2 treatments alter immunological synapse architecture and lytic granule polarization in primary NK cells. **a.** Representative images of primary NK cells isolated from PBMCs of three normal donors seeded on ICAM-1, anti-NKp30 and anti-NKp46, stained for F-actin (green), perforin granules (yellow) and nuclei (DAPI) and either untreated or treated with four concentrations of CK-869 or SMIFH2. Scale bars: 10 µm. **b-c** Graphs representing the fold changes of immunological synapse parameters of primary NK cells treated with (**b)** CK-869 and (**c)** SMIFH2 with respect to untreated controls. The data represent the mean of 4 replicates for each drug concentration (60-409 cells).

### Immunological synapse defect is associated with impaired cytotoxicity in CD8^+^ T cells from ARPC1B deficient patients

We then reasoned that the HCI approach might be adapted to characterizing lymphocyte impairments in the context of pathology. For that purpose, we implemented a morphological profiling of CD8^+^ T cells isolated from two patients suffering from a primary immunodeficiency caused by mutations in *ARPC1B,* which encodes one subunit of the ARP2/3 complex. Cells from the two patients and three normal donors were stimulated with ICAM-1 and either 1 or 10 µg/ml anti-CD3 Ab, and stained for perforin and LFA-1, F-actin and the nucleus. Examination of representative images suggests that the CD8^+^ T cells from both patients spread less than control cells and failed to assemble a typical IS (Fig. 5a). Analysis of 16 selected morphological features highlighted that F-actin intensity was decreased in T cells from the two patients, as compared to the cells from the normal donors, following stimulation with anti-CD3 Ab at both concentrations (Fig. 5b and **Fig. S3a**). This is comparable to the data collected above in cell lines and primary cells upon treatment with the ARP2/3 inhibitor CK-869. Cells from the two patients however displayed distinct morphological aberrations. While cell area was mostly affected in cells from patient 1, implying an impaired spreading ability, cell roundness was prominently decreased in patient 2, most likely resulting from aberrant peripheral actin spikes. T cells from patient 2 displayed an increased number and dispersion of perforin granules, similarly with what we observed in primary NK cells treated with CK-869. However, fewer perforin granules were detected in T cells from patient 1. Their dispersion was reduced in absolute terms but increased when normalized for the cell area, the latter being reduced. LFA-1 intensity was increased in the T cells from both patients and LFA-1 was localized at the cell rim rather than at the cell to substrate contact area, suggesting abnormal distribution of adhesive forces. Our morphological profiling clearly establishes that CD8^+^ T cells from the two considered ARPC1B-deficient patients have severe impairments in IS assembly. The distinct synaptic alterations revealed by our approach in the two patients could not be explained by differences in the phenotype of the cells, which showed similar expression of CD8, perforin, LFA-1, and granzyme B (Fig. S3 f-i). To further enrich our analysis, we applied once more a UMAP approach to explore the morphological space, which evidenced a marked segregation of the two patients from the control donors, but also among each other (Fig. 5c and **Fig. S3b**). This analysis therefore reinforces the finding that patient T cells have aberrant synaptic traits and that the nature of these aberrations is distinct between the two patients. Interestingly, we noticed that even though the normal donors clustered closely to each other, donor 3 did not overlap with the other two at either anti-CD3 Ab concentration, confirming that heterogeneity in IS morphology exists among normal donors, as observed above for NK cells. A first random forest model showed that we could determine the concentration of anti-CD3 Ab used to stimulate the cells with an accuracy of 95% and *F*_1_ score of 0.95 (**Fig. S3c**), and indicated that IS assembly varied according to the concentration of anti-CD3 Ab (**Fig. S3d**). This was associated with changes in shape and radial distribution in the cytoplasm, while cytoplasm intensities were the most discriminative feature category in average, fitting a scenario in which TCR stimulation strength would differentially remodel the actin cytoskeleton and associated synapse morphology. A second model showed that our approach is powerful enough to distinguish patient cells from normal donor cells seeded on ICAM-1 and 10 µg/ml anti-CD3 Ab by achieving a perfect classification on a validation set (Fig. 5d), distinguishing ARPC1B deficient cells mostly on the basis of textural and intensity distribution changes within the cytoplasm (Fig. 5e). Moreover, some features changed not only between ARPC1B patients and normal donors but were as well discriminating between patient 1 and patient 2 (**Fig. S3e**), further reinforcing that the two patients have distinct IS architectures. The aberrant IS characterized in the patients through the morphological profiling approach alluded to a possible functional defect. We therefore assessed the cytotoxic activity of CD8^+^ T cells towards anti-CD3 Ab-coated P815 target cells. Whereas, normal CD8^+^ T cells started to kill target cells in four hours, CD8^+^ T cells from the ARPC1B deficient patients failed to do so. The patient derived CD8^+^ T cells remained defective at killing target cells over a prolonged 24-hour incubation (Fig. 5f). This indicates that the T cells from the patients most likely fail to secrete lytic molecules, despite a normal content in perforin and granzyme B (**Fig. S3g** and **i**). Our results therefore indicate that the defects in IS organization characterized in both patients are leading to a severe impairment of the cytotoxic activity.

**Fig. 5.**
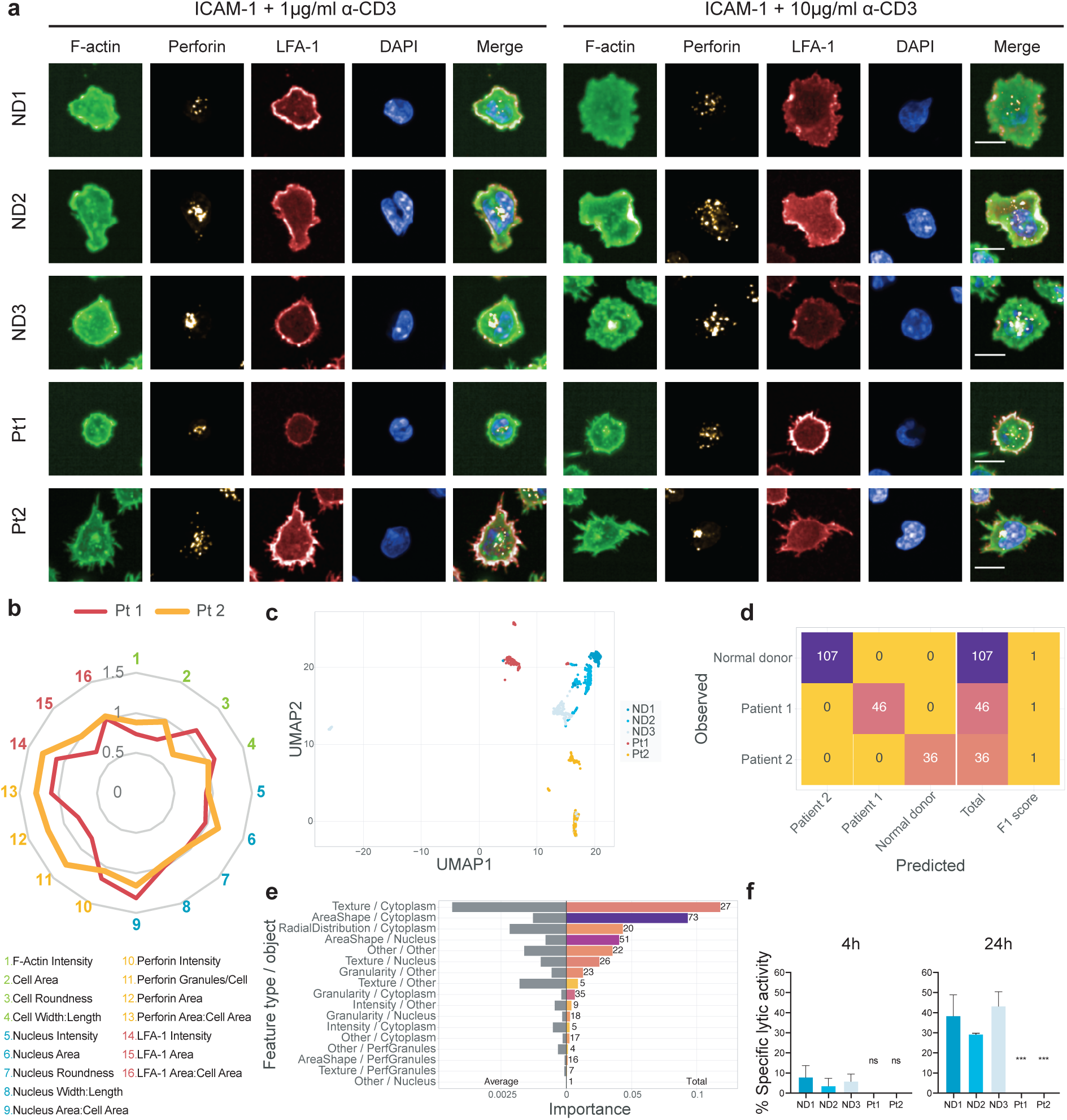
High content imaging of the immunological synapse in ARPC1B deficient CD8+ T cells. **a.** Representative images of CD8+ T cells from normal donors and ARPC1B deficient patients seeded on ICAM-1 and either 1 or 10 µg/ml anti-CD3 and stained for F-actin (green), perforin granules (yellow), LFA-1 (red) and nuclei (DAPI). Scale bars: 10 µm. **b.** Characteristics of the immunological synapse of CD8+ T cells of the two ARPC1B deficient patients represented as fold change with respect to the average of the three normal donors seeded on ICAM-1 and 1 µg/ml anti-CD3. The data represents the mean of 6 replicates for each donor (10687-19353 cells) **c.** UMAP projection of CD8+ T cells morphological profiles from the two patients and the three normal donors seeded on ICAM-1 and10 µg/ml anti-CD3. **d.** Confusion matrix and class-wise performance on held-out data of a random forest model trained to discriminate between patient and normal donors based on the morphology of CD8+ T cells seeded on ICAM-1 and 10 µg/ml anti-CD3. **e.** Total and average importance of morphological features per measurement type and biological object described for the prediction of patient and normal donor CD8+ T cells seeded on ICAM-1 and 10 µg/ml anti-CD3. **f.** Specific lytic activity of patient and normal donor CD8+ T cells incubated with P815 cells coated with 10 µg/ml anti-CD3 after 4 h and 24 h. Values represent the mean of triplicates and error bars show SD. Significance is noted as *(P <0.05).

## Discussion

By combining automated cell imaging with computational image analysis pipelines, HCI provides novel opportunities to systematically analyze cellular mechanisms^15, 36, 37^. However, the potential of such approach has not yet been explored for the study of immune cells. We here tailor a HCI approach for the high-resolution morphological profiling of various human cytotoxic lymphocyte population, and focus on the imaging of the IS as a mean to capture the activation state and effector potential of these cells. We validate our HCI approach by identifying distinct morphological signatures evoked by a panel of actin-targeting drugs. We further reveal the power of our HCI approach to discriminate individual donors on the basis of immune cell morphological traits. We also exemplify the clinical applicability of such approach by identifying cytotoxic lymphocyte aberrations in patients with a severe congenital immunodeficiency.

Although we use a minimalistic 2D static approach based on the adsorption of stimulatory molecules on the surface microwells, it proves to robustly stimulate the assembly of morphological structures qualifying as IS. We show that various human lymphocyte populations, including model cell lines, cells freshly isolated from the blood, as well as expanded primary cells can be stained and imaged with an automated confocal microscope at high resolutive power in a 384-well format, allowing the analysis of several samples, activation conditions and perturbations in parallel. Computationally, we use robust statistics and work at an image-level resolution, typically gathering a few dozens of cells imaged over four z planes representing the 2-µm section of the cells most proximal to the stimulatory substrate. While most morphological profiling studies have been limited to average profiles over wells or replicates^14, 37, 38^, a few approaches have defined profiles based on single cells ^39, 40^. We here rather consider the variability in morphology displayed in each image by including measures of dispersion that are proven to be beneficial for morphological profiles and could potentially be further improved by adding a higher order joint statistical moment ^41^. From an analytical point of view, we elaborate two complementary methods. First, we focus on a pre-defined set of morphological features based on prior knowledge and including cell and nucleus shape parameters as well as intensities of F-actin, LFA-1 and perforin at the synaptic plane. We show that such method can be applied to relatively low numbers of images and provides meaningful identification of discriminative features when comparing experimental conditions. Second, we implement a high-resolution and unbiased morphological profiling pipeline, from which novel relevant features can be identified and from which high-performance classifiers can be trained to discriminate cell states corresponding to different stimulations, drug treatments or genetic defects.

Beyond the methodological advance provided in this study, we present data relevant to the understanding of lymphocyte activation in both a fundamental and medically relevant perspective. Among the pre-defined set of morphological features, we identify increase of F-actin as a hallmark of T and NK lymphocyte stimulation by combinations of ICAM-1 and antibodies directed against CD3 or NK receptors, respectively. This is in line with the previously established role of the actin cytoskeleton in driving the cell spreading behavior supporting IS assembly^42, 43^. The further investigation of the role of actin cytoskeleton remodeling by the treatment of NK cells with a drug array reveals distinct morphological alterations upon targeting actin polymerization, depolymerization and myosin II. Our data also point to converging morphologies induced by some of the drugs with distinct modes of action, possibly related to a limited number of configurations of the cytoskeleton, as recently described in an adherent neuroblastoma cell line^36, 44^. Strikingly, most tested drugs yield prominent alteration of the distribution of perforin-containing granules, indicating that the different facets of actin cytoskeleton dynamics are all important to regulate the polarized delivery of lytic granules at the IS ^25, 45^.

Owing to the distinct morphological profiles observed for each drug, and the detection of dose-dependent effects, both in cell lines and primary cells, such an approach could be applied in the context of immunotherapeutic drugs. A striking finding of the application of morphological profiling to lymphocyte populations is that it reveals a previously unappreciated level of heterogeneity in cellular morphological traits among individuals. When considering the data pertaining to the NK cells freshly isolated from the blood, we cannot rule out that morphological differences arise from distinct activation states of the cells from different donors. However, in vitro stimulation and expansion of T lymphocytes, which is expected to robustly drive cells towards a differentiated phenotype^46^, was also associated with distinct morphological traits. Further analysis of larger cohorts of donors and sorted subpopulations of lymphocytes will be required to precisely appreciate the degree of morphological heterogeneity among individuals and lymphocyte subsets. The detection of distinct morphological profiles among healthy individuals certainly highlights the extreme sensitivity of the HCI approach to characterize and compare cell populations. A further illustration of this property is provided by the characterization of IS alterations in T lymphocyte populations isolated from 2 patients with ARPC1B deficiency. Interestingly, again, our approach points to distinct morphological alterations in the cells from the 2 patients considered. Such differences might be inherent to the severity of the ARPC1B genetic defect. ^47, 48^ The study of larger cohorts of patients, which would be compatible with the herein developed approach, would be required to answer such question.

CD8^+^ T cells from ARPC1B patients display an aberrant IS morphology including defects pertaining to the distribution of perforin granules and LFA-1. Comparably to other studies, we show a reduced cell area and a failure to spread radially and emit lamellipodia upon TCR stimulation^47^. The lack of lamellipodia formation was also observed upon NK-92 and primary NK cell treatment with CK-869. Our data reveals increased accumulation of lytic granules at the IS for one patient, which could indicate a defect in granule exocytosis, opposed to a decrease in perforin related parameters for the other patient, most likely indicative of failed lytic granule polarization. These observations are in agreement with a recent study showing defective lytic granule polarization and degranulation in ARPC1B deficient CD8^+^ T cells^49^. The detection of such IS defects is suggestive of a possible alteration of the cytotoxic activity. Our data shows that ARPC1B deficient cells fail to eliminate target cells, as recently reported^49^. This illustrates the potential of HCI to provide guidance for the implantation of complementary low throughput assays to assess defects at the functional and molecular level. At this stage, we cannot generalize the case of the ARPC1B deficiency in establishing a systematic relationship between IS alteration and functional defect. However, it is interesting to mention that multiple primary immunodeficiencies have been found by us and others to associate IS defects and functional impairments^50–53^. Previous reports have also shown that PIDs where the IS is defective fail to eliminate target cells^51, 53, 54^ The systematic analysis of multiple such pathologies and corresponding cellular models would certainly provide a unique opportunity to establish rules linking morphology to function.

Overall, we provide here an innovative HCI approach to unbiasedly interrogate the biology of lymphocyte populations. It provides a rich way to identify and interpret details of the IS architecture and surpass current approaches in detecting morphological traits of specific lymphocyte populations, as illustrated by the distinct morphological profiles identified among the primary lymphocytes of individual donors. This hold promises to stratify patients based on specific morphotypes of lymphocytes or other leukocytes. Therefore, we thoroughly report both the experimental and computational methods employed and provide all scripts used in the analysis to maximize the reproducibility of the approach developed herein. We hope this encourages further research leveraging the application of HCI to blood-derived cell subsets, for potential translation in the field of cancer therapy and personalized medicine.

## Materials and methods

### Cell lines and primary cells

Jurkat cells were cultured in RPMI (Gibco) supplemented with 10% FBS, 1% penicillin/streptomycin, 1% sodium pyruvate, 1% non-essential amino acids and 1% HEPES (all from Thermo Fisher Scientific). NK-92 cells were cultured according to the recommendations from ATCC. Primary NK cells were purified from freshly-isolated PBMCs using the MagniSort Human NK enrichment kit (Invitrogen) and maintained in RPMI supplemented with 5% human serum, 1% penicillin/streptomycin, 1% sodium pyruvate, 1% non-essential amino acids and 1% HEPES. Primary CD8^+^ T cells were purified from frozen PBMCs of 3 healthy donors and 2 ARPC1B deficient patients by negative selection using the EasySep Human CD8^+^ T cell enrichment kit. CD8^+^ T cells were stimulated in RPMI supplemented with 5% human serum, 1% penicillin/streptomycin, 1% sodium pyruvate, 1% non-essential amino acids, 1% HEPES 1 µg/ml PHA and 100 IU/ml IL-2. CD8^+^ T cells were expanded for further rounds every 2 weeks with a mixture of irradiated PBMCs from 3 normal donors. Peripheral blood from healthy donors and patients was obtained in accordance with the 1964 Helsinki declaration and its later amendments or ethical standards. Informed consents were approved by the relevant local Institutional Ethical Committees.

### Culture and staining conditions used for High content imaging

CellCarrier Ultra tissue culture treated plates (Perkin Elmer) were coated with either 0.01% PLL (Merck) or a combination of 2 µg/ml ICAM-1 (R&D Systems), 1 µg/ml NKp30 (R&D systems, MAB18491) and 1 µg/ml NKp46 (BD Biosciences, 557487). NK-92 cells were cultured in IL-2 free medium overnight. 15000 NK-92 and 5000 primary NK cells were seeded per well and left for 30 min at 37°C to adhere and form the synapse. Cells were fixed with 3% paraformaldehyde (Merck) and stained with anti-perforin Ab and phalloidin-AF 488. Goat anti-mouse AF 555 was used to reveal perforin staining. Nuclei were stained with DAPI.

NK-92 were treated with 5, 10 and 50 µM Blebbistatin, 10, 25 and 50 µM CK-869 (Merck), 0.1, 1 and 2.5 µM Jasplakinolide (Merck), 0.1, 0.25 and 0.5 µM Latrunculin B (Merck), 50, 100 and 250 µM SMIFH2 (Merck), 10 50 and 100 µM Wiskostatin (Merck) and 5, 10 and 25 µM Y-27632 (Merck) for 30 min at 37°C and washed twice in PBS before seeding onto the plates and letting them adhere for 30 min. The same procedure was applied to primary NK cells treated with 5, 10, 25 and 50 µM CK-869 and 25 50, 100 and 250 µM SMIFH2.

CellCarrier Ultra multiwell tissue culture treated plates were coated with either 0.01% poly-L-lysine or a combination of 2 µg/ml ICAM-1 and 10 µg/ml anti-CD3 (eBioscience). 10000 Jurkat or 5000 CD8^+^ T cells were seeded per well and left for 15 min at 37°C to adhere and form the synapse. Cells were fixed with 3% paraformaldehyde and stained with anti-LFA-1 (BioLegend, 301202) and phalloidin-AF 488 (Thermo Fisher Scientific) in permeabilization buffer (eBioscience). Goat anti-mouse AF 647 antibody (Thermo Fisher Scientific, A-21240) was used to reveal LFA-1 staining. CD8^+^ T cells were in addition stained with anti-perforin and goat anti-mouse AF 555 (Life technologies) was used to reveal perforin staining. Nuclei were stained with DAPI (Thermo Fisher Scientific). Stained cells were kept in PBS at 4°C until imaging.

### Image acquisition and processing

Images were acquired on an automated spinning disk confocal HCS device (Opera Phenix, Perkin Elmer) equipped with a 40x 1.1 NA Plan Apochromat water immersion objective and a sCMOS camera. For each well, 40 randomly selected fields and 8 stacks per field (0.5 µm step) were acquired. Stacks of images were combined with maximum projection for four focal slices (*z* from 1 to 4 with a 0.5 µm step), then assembled in sets of images per field of view corresponding to DAPI, phalloidin and LFA-1 or perforin depending on the cell type imaged. These datasets were processed, and measurements were made using CellProfiler 3.0^18^ (see Supplementary Files [CellProfiler pipeline]). In brief, we assess the image quality, log-transform the intensities for experiments with high background noise, correct the illumination on each image based on background intensities, avoid DNA precipitations by multiplying intensities on DAPI channel by phalloidin intensities before segmenting cell nuclei using global minimum cross entropy thresholding. We perform a secondary segmentation of the cytoplasms using the watershed method^55^ and global minimum cross entropy thresholding on the phalloidin channel. Image sets with low maximal DNA intensity or showing no nucleus were discarded. Cells having more than 30% of their cytoplasm surface at less than 5 pixels of another cell were removed, in order to ignore clusters of cells and to focus on single cells displaying an IS. We segmented small actin speckles in the cytoplasm at more than 3 pixels from the membrane as well as speckles of perforin and secondary objects spanned around the nuclei by LFA-1 staining. Additionally, primary NK and expanded CD8^+^ T cells associated with less than 2 perforin granules were excluded from the analysis. Finally, we measured colocalization of these objects, intensities in the nuclei and cytoplasms, granularity on all channels, textural and shape features, intensity distributions, distance and overlap between objects and counted speckles neighbors less than 10 pixels away. We then kept the average and the standard deviation of these features per field of view. This led to 1898 and 2076 morphological features in NK92 and Jurkat cells respectively. For primary NK cells and expanded CD8^+^ T cells, features related to actin speckles were excluded, as they were not found to be informative, resulting in 2386 and 1517 features, respectively.

Stacks of images were combined with maximum projection for four focal slices (*z* from 1 to 4 with a 0.5 µm step), then assembled in sets of images per field of view corresponding to DAPI, phalloidin and LFA-1 or perforin depending on the cell type imaged. These datasets were processed, and measurements were made using CellProfiler 3.0^18^ (see the pipelines provided).

In brief, we assess the image quality, log-transform the intensities for experiments with high background noise, correct the illumination on each image based on background intensities, avoid DNA precipitations by multiplying intensities on DAPI channel by phalloidin intensities before segmenting cell nuclei using global minimum cross entropy thresholding. We perform a secondary segmentation of the cytoplasms using the watershed method^55^ and global minimum cross entropy thresholding on the phalloidin channel. Image sets with low maximal DNA intensity or showing no nucleus were discarded. Cells having more than 30% of their cytoplasm surface at less than 5 pixels of another cell were removed, in order to ignore clusters of cells and to focus on single cells displaying an IS. We segmented small actin speckles in the cytoplasm at more than 3 pixels from the membrane as well as speckles of perforin and secondary objects spanned around the nuclei by LFA-1 staining. Additionally, primary NK and expanded CD8^+^ T cells associated with less than 2 perforin granules were excluded from the analysis. Finally, we measured colocalization of these objects, intensities in the nuclei and cytoplasms, granularity on all channels, textural and shape features, intensity distributions, distance and overlap between objects and counted speckles neighbors less than 10 pixels away. We then kept the average and the standard deviation of these features per field of view. This led to 1898 and 2076 morphological features in NK92 and Jurkat cells respectively. For primary NK cells and expanded CD8^+^ T cells, features related to actin speckles were excluded, as they were not found to be informative, resulting in 2386 and 1517 features, respectively.

### Data processing and visualization

We subsequently conducted analyses in R 3.5.1 with the data visualization package ggplot2 3.1.1 and Microsoft Excel (Version 1902). We further selected a smaller set of informative morphological features and checked the quality of processed images by (i) removing wells with low maximal DNA intensity and cell count, (ii) removing features and images generating missing values and (iii) removing constant features in the study dataset or the subset of negative controls used as reference.

From these images passing our quality checks, up to 16 raw summary variables were extracted based on their interpretability and on their known relevance to describe ISs. The fold changes compared to unstimulated or untreated controls were further reported and displayed in the form or radar charts.

On the other hand, for all features, per-image values *X* were transformed successively with the following functions *f*_1_ and *f*_2_, with *X_Control_* the negative controls in *X* on which the data is normalized:

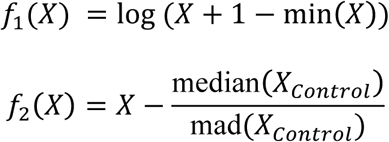

To remove redundancy in the set of features used for downstream analyses, we ensured that the selected variables were not excessively linearly correlated. To do so, all features were ordered from highest to lowest median absolute deviation (hence by variation in the experiment compared to negative controls). Starting from the top of this list, all other features linearly correlated to the first feature with a Pearson’s coefficient higher than 0.6 were excluded. We sequentially went on with the next remaining feature in the list and iterated until the acquisition of a small and informative set of uncorrelated features.

This set of features was used for visualization and quantification of the overall morphological changes induced by perturbations. We then reduced the dimensionality of the data using the UMAP algorithm^56^ to 2 dimensions for visualizations and 3 dimensions for computation of the statistical significance of morphological effects in the drug screen on NK92. This pipeline succeeded in selecting a wide range of features that were not excessively biased by confounders (**Fig. S2d**).

### Robust Morphological Perturbation Value

To quantify the significance of overall changes in morphology between a perturbed state and a reference state (healthy or untreated cells), we define the Robust Morphological Perturbation Value. This extends the concept of Multidimensional Perturbation Value^33^, which defines a single value summarizing the statistical significance of morphological changes in multidimensional spaces, by using robust statistics and the minimum covariance determinant^34^ decreasing the sensitivity to technical (unfiltered artifacts) and biological outliers (images displaying extreme morphologies or uncommon cell states). In brief, the RMPV is obtained for *X* the set of all filtered and uncorrelated features and *X_WT_* the subset of the data corresponding to images of the reference population in five steps. First, the minimum covariance determinant estimator *M*(*X_WT_*) is calculated to describe the variation of morphologies observed in the reference set, using its implementation in the R package *robustBase* version 0.93. Second, this value is used to determine *R*, the robust Mahalanobis distance of each images of *X* to *X_WT_* [arXiv:1904.02596 [stat.ME]]. Third, the median value *R̃* = median(*R*) was obtained for each drug tested. Fourth, for 2000 iterations the labels of the condition and the reference were randomly permuted to obtain an empirical distribution of *R̃* under the assumption that there was no difference between the multivariate location and scatter of the morphological parameters of the perturbation and the reference. Finally, the RMPV is defined as the empirical p-value obtained from these distributions after FDR adjustment for testing changes in multiple conditions and indicates the probability of observing at least half of the images displaying morphological changes of a similar intensity if the perturbation was similar to the reference.

### Random forest classification and interpretation

Using the set of informative and uncorrelated morphological features – previously used for dimensionality reduction, we trained a random forest classifier^57^ using the R package *randomForest* version 4.6.

Using the set of informative and uncorrelated morphological features – previously used for dimensionality reduction, we trained a random forest classifier^57^ using the R package *randomForest* version 4.6. Each forest included 1000 decision trees. The data was split in 6 folds of equal size, each containing all possible classification label. To select the optimal number *mtry* of variables selected at each split, we incremented the parameter value from 20 to 90 by steps of 10 and assessed the performance using the macro *F*_1_ score as defined below in a 5-fold cross-validation scheme. One extra fold was used as validation set to estimate the performance of the model after selection of the optimal parameters and retraining on all of the 5 folds used for cross-validation. In the case of the drug screen on the NK-92 cell line, we used a similar approach using the 13 features of known relevance in describing the IS as input, and testing *mtry* values from 1 to 13 with steps of 3. Overall the performance was evaluated using the macro *F*_1_ score:

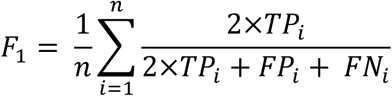

where *n* is the number of categories in the classification, and *TP_i_*, *FP_i_* and *FN_i_* are respectively the number of true positives, false positives and false negatives for category *i* in the validation set. To interpret the feature importance in the prediction, we extracted the mean decrease in accuracy obtained when including each feature, either for the prediction of a given class or overall using micro averaging. The total and average importance of features split in distinct groups based on the type of measurements and biological object described were calculated as well. These feature groups were defined based on the corresponding CellProfiler measurement types and biological objects. Features that did not describe the cytoplasm, nucleus, perforin granules or actin granules were counted in the “Other” biological object category. Similarly, features that did not correspond to the “Texture”, “AreaShape”, “RadialDistribution”, “Granularity” or “Intensity” measurements were grouped under the term “Other”.

### Cytotoxicity assay

Target P815 cells were stained for 30 min with Cell Tracker green CMFDA (Thermo Fisher Scientific) and coated with 10 µg/ml anti-CD3 (eBiosciences, 16-0037-81) for one hour at 37°C. They were also treated with 0.2 µg/ml of aphidicolin to prevent their proliferation. P815 were incubated with effector CD8^+^ T cells in U-bottom 96 well plates at an effector: target ratio of 1:1 for 4 and 24 hours^58^. Cells were then stained with Amino-Actinomycin D (7-AAD) (BD Biosciences) to discriminate dead and alive cells using the MacsQuant VYB (Miltenyi) and analyzed with FlowJo. The number of residual alive CMFDA^+^ / 7-AAD^-^ cells was assessed to calculate cytotoxicity. Student’s *t*-test was used to calculate significance.

### Phenotypic analysis

Expanded CD8^+^ T cells from normal donors and ARPC1B-deficient patients were stained with fluorochrome-coupled antibodies recognizing the extracellular markers CD8 (BioLegend, 344718) and LFA-1 (BioLegend, 363404) for 30 min at 4°C. Intracellular staining was performed following fixation and permeabilization, with the following antibodies perforin (BioLegend, 308110) and granzyme B (BDPharmigen, 561142) for 45 min at 4°C. The data were acquired on MacsQuant Q10 (Miltenyi) and analyzed with FlowJo. Student’s *t*-test was used to calculate significance.

### Data availability

All the CellProfiler pipelines and morphological measurements used in this analysis are made available on FigShare with the DOI 10.6084/m9.figshare.11619960 [already available for reviewers with the following private link: https://figshare.com/s/3c06753839d77783a899].

### Code availability

The analyses can be found and reproduced using the Docker image and scripts provided on Github and identified with the DOI 10.5281/zenodo.3518233.

## Acknowledgements

The authors thank Fatima-Ezzahra L’Faqihi-Olive, Valérie Duplan-Eche, Anne-Laure Iscache and Lydia De La Fuente-Vizuete from the cytometry platform of the CPTP and Isabelle Fernandes from the Organoid platform of the IRSD, Muriel Quaranta-Nicaise, Laurène Pfajfer, Michael Caldera, Marianne Guisset and Anton Kamnev for discussion and technical advice, Alessandro Aiuti, Marco Gattorno and Stefano Volpi for patient-derived cell lines. Y.G. performed the experiments, analyzed the results and wrote the paper, L.V. designed and performed data analysis and wrote the paper. A.R. contributed in image acquisition. A.F., J.M. and K.B. participated in research design and scientific discussions. L.D. designed the research, supervised the experiment analysis and wrote the paper.

The authors declare no competing interests.

