## Supplementary material for "Morphological profiling of human T and NK lymphocytes identifies actin-mediated control of the immunological synapse": Suppl figures S1-S3 and Tables

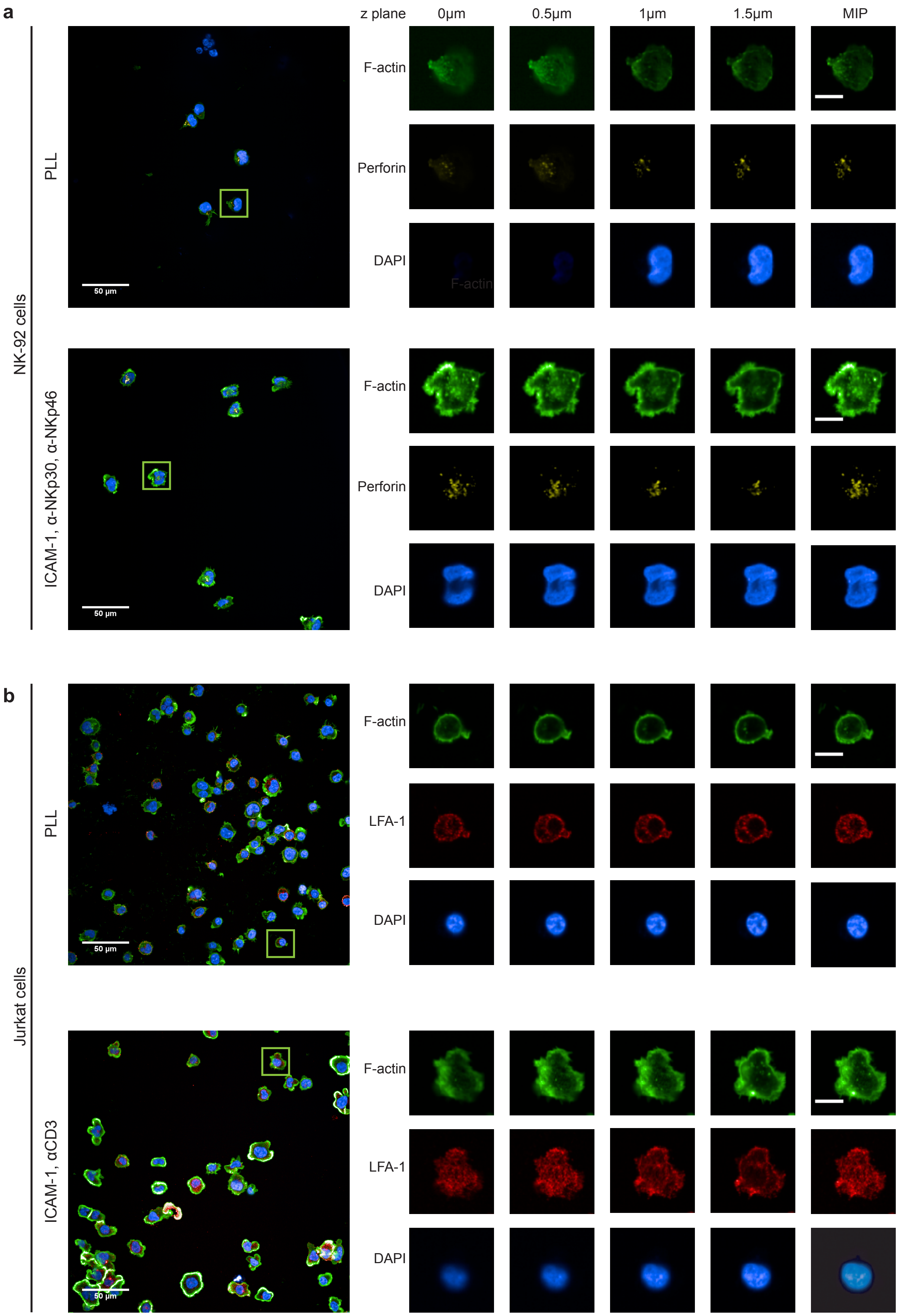

a
b
z plane
0µm
0.5µm
1µm
1.5µm
MIP
NK-92 cells
PLL
ICAM-1, α-NKp30, α-NKp46
F-actin
Perforin
DAPI
50 µm
Jurkat cells
ICAM-1, αCD3
LFA-1

**Supplementary Fig. 1 | Individual channels and z planes of single cells isolated from representative fields of view.** a. Maximum intensity projection (MIP) of a representative field of view of NK-92 cells seeded on PLL (top) or ICAM-1, anti-NKp30 and NKp-46 (bottom), with zoom on a single representative cell stained for F-actin (green), perforin granules (yellow) and nuclei (DAPI) imaged at 4 z-planes with a step of 0.5 μm and its MIP. Scale bars: Field of view 50 μm and single cell 10 μm. b. MIP of a representative field of view of Jurkat cells seeded on PLL (top) ICAM-1, anti-CD3 (bottom), with zoom on a single cell stained for F-actin (green), LFA-1 (red) and nuclei (DAPI) and imaged at 4 z-planes with a step of 0.5 μm and its MIP. Scale bars: Field of view 50 μm and single cell 10 μm.

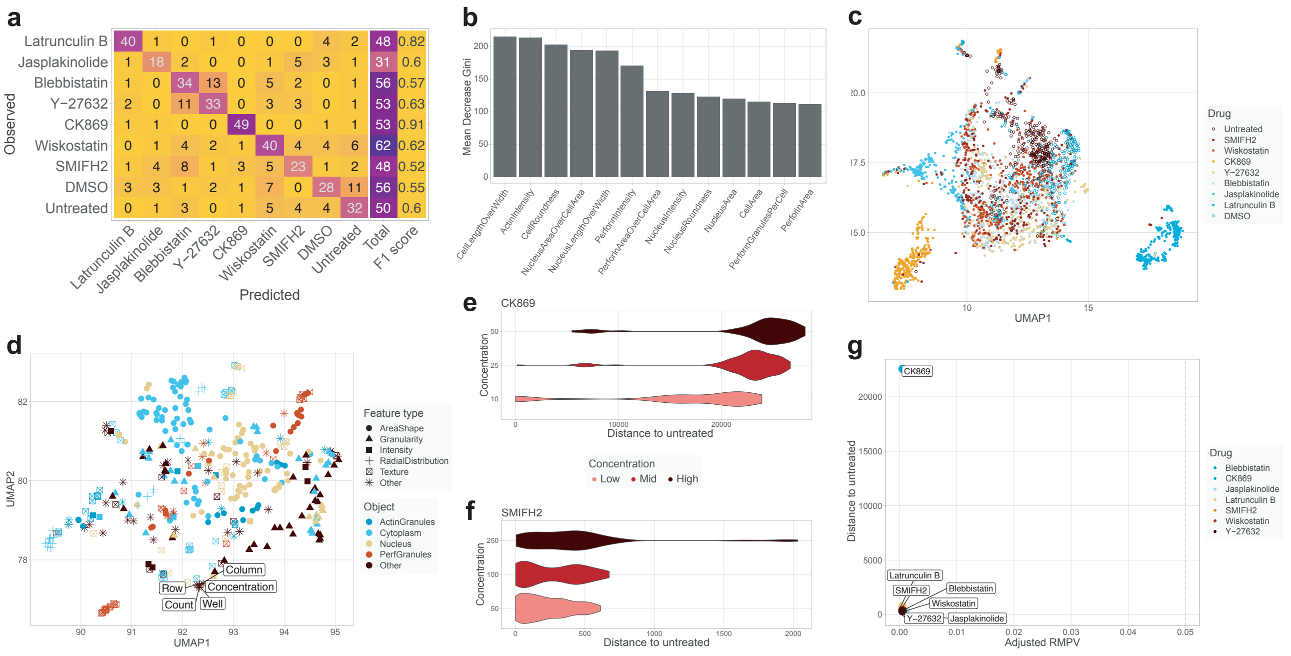


**Supplementary Fig. 2 | Drug treatment leads to distinct immunological synapse phenotypes in NK-92 cells.** a. Confusion matrix and class-wise performance of a random forest model trained to predict drug treatment based on 13 hand-picked morphological features of NK-92 seeded on ICAM-1, anti-NKp30 and anti-NKp46. b. Importance of the 13 morphological parameters for the classification described in panel (a). c. UMAP representing the clustering of all the drugs and the untreated conditions. d. UMAP representing the relations between confounders and morphological features, obtained by fitting the UMAP on the transpose of the data underlying panel (c). e-f. Violin plots representing the effect size of drug concentrations on morphological features for (e) CK-869 and (f) SMIFH2. g. FDR-corrected Robust Morphological Perturbation Value (RMPV) of the different drugs.

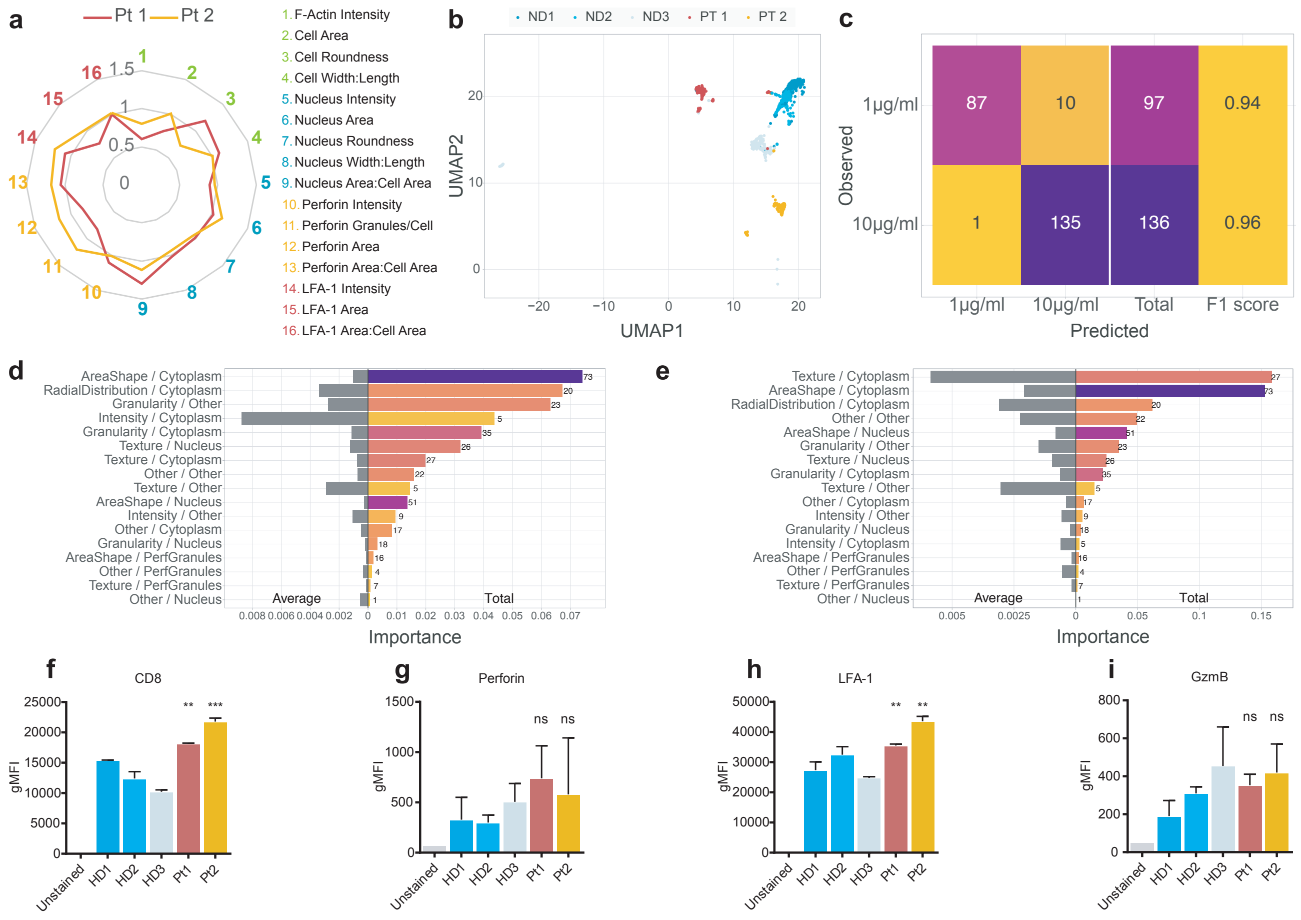


**Supplementary Fig. 3 | Distinct immunological synapse phenotype in CD8+ T cells from ARPC1B-deficient patients.** a. Characteristics of the immunological synapse of the CD8+ T cells of the two ARPC1B deficient patients represented as fold change with respect to the average of the three normal donors seeded on ICAM-1 and 1 µg/ml anti-CD3. The data represents the mean of 6 replicates for each donor (7525-17725 cells). b. UMAP of cells of the ARPC1B patients and the normal donors seeded on ICAM-1 and 1 µg/ml anti-CD3. c. Confusion matrix and class-wise performance of a random forest model trained to predict the concentration of anti-CD3 antibodies based on the morphology of normal donors lymphocytes stimulated with ICAM-1 and either 1 or 10 µg/ml anti-CD3. d-e. Total and average importance of measurement type and biological object described in the prediction of whether images corresponded to (d) normal donors CD8+ T lymphocytes seeded on ICAM-1 and 1 or 10 µg/ml anti-CD3 or (e) to normal donors, patient 1 or patient 2 lymphocytes stimulated with ICAM-1 and 10 µg/ml anti-CD3. f-i Phenotyping of expanded CD8+ T cells of ARPC1B patients and normal donors showing mean fluorescence intensity of (f) CD8, and (g) perforin, (h) LFA-1, and (i) Granzyme B in the CD8+ population. Values represent the mean of duplicates and error bars show SD. Significance is noted as ns (P >0.05), ** (P<0.01) and *** (P<0.001).

| | NK-92 | | |
|---|---|---|---|
| | PLL | ICAM-1, α-NKp30, α-NKp46 | Fold increase |
| F-Actin intensity | 0.065098277 | 0.110223848 | 1.69319149 |
| Cell Area ($\mu m^2$) | 2469.321237 | 3130.618434 | 1.26780525 |
| Cell Roundness | 0.674079624 | 0.444435641 | 0.65932217 |
| Cell Width:Length | 0.686074044 | 0.71563093 | 1.04308119 |
| Nucleus Intensity | 0.067203031 | 0.067329026 | 1.00187485 |
| Nucleus Area(μm2) | 1037.718811 | 1333.179255 | 1.2847211 |
| Nucleus Roundness | 0.706061099 | 0.689874902 | 0.97707536 |
| Nucleus Width:Length | 0.685545171 | 0.706542199 | 1.03062822 |
| Nucleus Area:Cell Area | 0.427455169 | 0.431859895 | 1.01030453 |
| Perforin Intensity | 0.018403047 | 0.018502314 | 1.00539403 |
| Perforin Granules/Cell | 13.04941437 | 16.7196273 | 1.28125499 |
| Perforin Area(μm2) | 86.05831253 | 117.5449158 | 1.36587521 |
| Perforin Area:Cell Area | 0.035039484 | 0.036882184 | 1.05258922 |

**Supplementary Table 1 | Mean values and fold increase of immunological synapse parameters in NK-92 cells.** Mean values of individual parameters pertaining to the immunological synapse in NK-92 cells seeded on PLL or ICAM-1, anti-NKp30 and anti-NKp46, and the fold change of the ratio of each mean value on the stimulated condition with respect to PLL. Intensity is measured in arbitrary units and area in $\mu m^2$.

| | Jurkat | | |
|---|---|---|---|
| | PLL | ICAM-1, α-CD3 | Fold increase |
| F-Actin intensity | 0.000289437 | 0.000437181 | 1.5104544 |
| Cell Area(μm2) | 1921.009837 | 2690.327023 | 1.4004754 |
| Cell Roundness | 0.530654443 | 0.61184158 | 1.15299436 |
| Cell Width:Length | 0.800192404 | 0.767932394 | 0.95968468 |
| Nucleus Intensity | 0.07313481 | 0.074429279 | 1.01769975 |
| Nucleus Area(μm2) | 879.0221429 | 995.945146 | 1.13301486 |
| Nucleus Roundness | 0.830340643 | 0.806271514 | 0.97101295 |
| Nucleus Width:Length | 0.779080494 | 0.754652368 | 0.96864493 |
| Nucleus Area:Cell Area | 0.460560432 | 0.379756314 | 0.82455263 |
| LFA-1 Intensity | 0.006679646 | 0.007880989 | 1.17985131 |
| LFA-1 Area(μm2) | 1728.306565 | 2526.153065 | 1.46163483 |
| LFA-1 Area:Cell Area | 0.900646361 | 0.938478533 | 1.04200558 |

**Supplementary Table 2 | Mean values and fold increase of immunological synapse parameters in Jurkat cells.** Mean values of individual parameters pertaining to the immunological synapse in Jurkat cells seeded on PLL or ICAM-1 and anti-CD3, and the fold change of the ratio of each mean value on the stimulated condition with respect to PLL. Intensity is measured in arbitrary units and area in $\mu m^2$.
